# Remodeling of the hepatic circadian transcriptome across the estrous cycle

**DOI:** 10.64898/2026.06.22.733912

**Authors:** Kiandra A. Smith, Tsedey Mekbib, Aili M. Fisher, Aisha Rollins-Hairston, Ting-Chung Suen, Hao A. Duong, Morris Benveniste, Jason P. DeBruyne

**Author notes:** Kimberly-Clark Inc, Roswell GA 30076. **Corresponding Author information:** Jason P. DeBruyne, Department of Pharmacology and Toxicology, Morehouse School of Medicine, 720 Westview Dr., Atlanta GA, 30310. **Competing Interest Statement:** None.

## Abstract

The circadian clock system drives rhythms in gene expression, tailoring an organism’s behavior and physiology to the ∼24-hour day-night environmental cycle. The mechanisms underlying this system are believed to be largely the same between adult males and females, but recent findings are starting to challenge this notion. Menstrual/estrous cycles (e-cycles) in females modulate a variety of circadian-controlled behaviors, but how they may interact with rhythmicity at the system level are an unknown but vital dimension in circadian biology in females. To start bridging this gap, we explored the circadian transcriptome of the mouse liver across the four phases of the ‘free-running’ e-cycle. Much to our surprise, we found that the e-cycle imposes a∼4-day rhythm in the 24-hour rhythmicity of a considerable proportion of the circadian transcriptome, with the largest differences in rhythmicity occurring between pre- and post-ovulation phases. These differences also appear highly time-of-day dependent, and gene promotor analysis suggests that both e-cycle and sex dependent rhythmicity may be due to relatively simplistic mechanisms, possibly involving the action of core circadian transcription factors on output genes. Taken together, this study establishes that the estrous cycle modulates circadian gene expression in the liver, suggesting that circadian system has evolved specific mechanisms tailored to the different needs of males and females in their normal daily environments.

## Introduction

Circadian clocks are ubiquitous and drive daily gene expression rhythms in most tissues, which tailor an organism’s physiologies and behaviors to environmental day-night cycles. Circadian timekeeping is encoded by rhythmic “clock gene” expression, which is the product of a precise transcriptional, post-transcriptional, and post-translational negative feedback loop system comprised of at least three interconnected loops(1–3). These loops also drive rhythmic expression of 1,000’s of ‘output’ genes through either direct interaction with their promoters, or by interactions with other transcription factors. The subsequent rhythms in ‘output’ gene expression ultimately produce overt circadian rhythms in physiology and behavior.

Most circadian studies have been male-centric but have been extraordinary in dissecting the circadian clockwork mechanisms. While it has long been observed that several aspects of circadian rhythmicity display sex-specificity (4–9), underlying circadian timing mechanisms so far appear to be similar between males and females. However, both the rhythms driven by the clock and how the clock responds to environmental challenges are influenced by sex (9). Notably, rhythmicity across the transcriptome appears to be surprisingly different between males and females, but the mechanisms underlying why some genes are rhythmic only in females, for example, are only starting to be elucidated (6,10,11). The identification of female-specific circadian regulators such as SIAH2, insinuate that there may be yet-to-be discovered sex-dependent circadian mechanisms (12). Identification of these mechanisms is essential as all of these findings suggest that the lessons learned from male-centric studies may not broadly apply to females, and vice versa.

The menstrual/estrous cycle (e-cycle) in adult females also interacts with many of the same physiological rhythms driven by the circadian clock. In most rodents, the e-cycle has a period of ∼4-5 days and is coordinated by a hormonal interaction network involving hypothalamus, ovary (estrogen, progesterone), and pituitary (LH, FSH; HPG axis). E-cycles depend on a functioning circadian clock, although the circadian clock functions in the absence of e-cycles. E-cycles influence circadian locomotor, feeding, and body temperature patterns (8), and the disruption of either circadian or estrous cycling disrupts metabolic homeostasis, often resulting obesity and metabolic syndrome. However, whether these phenotypes have a related origin or are converging from distinct mechanisms is unknown.

We have therefore started to explore this question by assessing circadian regulation of the transcriptome across the ‘free running’ estrous cycle in the mouse liver, a highly sexually dimorphic tissue intricately involved in metabolism. Our data suggest that the e-cycle imposes a ∼4-day rhythm in the circadian expression of a large proportion of rhythmic genes. This cyclical remodeling occurs predominantly in a time-of-day specific manner, suggestive of discreet mechanisms. Moreover, this remodeling appears to diversify overall sex differences in hepatic rhythmicity, revealing new potential areas in which e-cycle may modulate overall liver function.

## Results

### Core clock genes oscillate independently of e-cycle

E-cycles in mice typically last 4-5 days and can be erratic, but are divided into four stages that occur in sequence: diestrus (D), proestrus (P), estrus (E), and metestrus (M) (13). To evaluate the potential impact of the estrous cycling on daily transcriptomes, we harvested tissues at 2-hour intervals from female mice across each of the four e-cycle stages (∼1 day per stage; Fig 1A, see Methods). For this initial trial, we harvested tissues from 1 animal at 2-hour intervals across 4 days as this design provides an optimal intersection of robustness in rhythm detection (14,15) and animal usage while incorporating daily e-cycle staging. E-cycle staging was done using vaginal lavages (16,17) performed daily between ZT2-4 (see Methods) for ∼12 days before harvesting tissues in an attempt to establish a baseline estrous cycle rhythm for each animal (Fig S1). However, as anticipated, we found that individual e-cycling was highly variable and did not follow a forecastable 4-5 day ‘rhythm’ (16) (Fig S1). We therefore utilized serum estradiol and follicle-stimulating hormone (FSH) levels, measured at the time of collection, to corroborate e-cycle phases determined by vaginal cytology that same day (Fig 1B). Of the 69 mice we started with (Fig S1), we selected 48 mice in which serum hormone measures corroborated vaginal lavage staging stage for analysis of the liver transcriptomes (Fig 1A-C).

**Figure 1.**
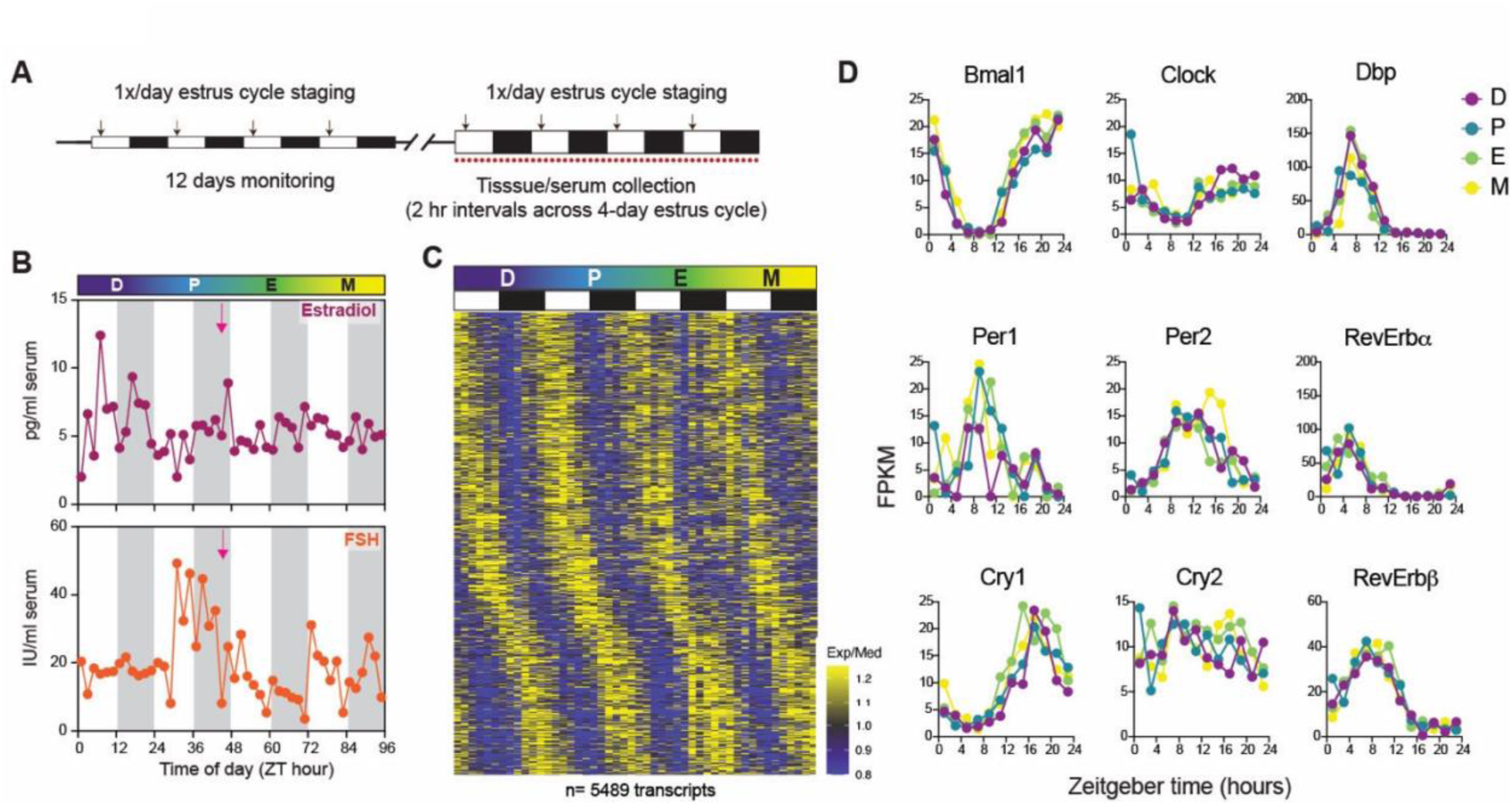
Female livers exhibit robust diurnal rhythms in gene expression across the estrous cycle without systematic effect on core clock gene expression. **A.** Experimental paradigm of e-cycle circadian collection. Alternating white/black bars represent the light-dark (LD) cycle, arrows indicate the timing of vaginal swabbing (ZT2-4), the red dots indicate 2-hr intervals for tissue/serum collection. A single mouse was collected at each 2-hour interval for each of the 4 e-cycle phases. **B.** Serum hormone levels of mice whose livers were subjected to transcriptome analyses. The LD cycle is indicated by the white/grey shading, and the pink arrow indicates the expected time of ovulation. **C.** Heatmap of rhythmic genes (see Fig S3), organized by e-cycle phase (D=diestrus; P= proestrus; E=estrus; M=metestrus) and time-of-day of peak expression (white/black bars indicate the 12:12LD cycle). **D.** Diurnal expression profiles of core clock genes at each estrous cycle phase.

We first compiled the transcriptome across the 96-hour collection period to examine potential e-cycle responsive genes and overall circadian dynamics (Dataset S1). Surprisingly, we found relatively small numbers of transcripts (<1%) that cycled in abundance with a ∼96-hour period. Similarly, we found that expression of relatively few transcripts (<2%) correlated with serum hormone levels (max r = 0.55, Pearson Correlation; Fig S2). In contrast, we found that ∼20% of detected transcripts appear to cycle with a ∼24-hour period across the 4-day collection (Fig 1C). Of note, with exception to a subtle but significant alteration in *Bmal1* expression, the expression of most genes within the circadian feedback loops did not appear to vary with e-cycle phase (Fig 1D, see also below). Combined, these data suggest that e-cycle and related hormones, across the dynamic physiological range, may have a relatively modest direct influence on hepatic gene expression or circadian clock function, and imply that circadian cycles may play a broader role in regulating liver function than the 4-day estrous cycle (18,19).

### Circadian transcriptome varies substantially with the e-cycle

To uncover changes in circadian gene expression patterns across the e-cycle (Dataset S1) while minimizing the potential impact of outliers since we collected single animal per timepoint, we utilized a moving average concept in which the rhythmic expression was defined using data from 2 full circadian cycles (24 animals), in rolling 48-hour intervals (Fig 2A). This approach improves reliability of detecting rhythmic transcripts compared to analyzing individual 24-hour periods for circadian-like cycling, as well as improves robustness for measuring rhythm phase and amplitude of individual genes (14). We also employed JTK-cycle to identify “rhythmic” transcripts and quantify rhythm phase and amplitude, as this method is less sensitive to the influence of outliers (see Methods) (15,20). However, we note a limitation of the rolling two-day approach is that it may not be able to reliably detect rhythmicity that occurs in a single e-cycle stage/day.

**Figure 2.**
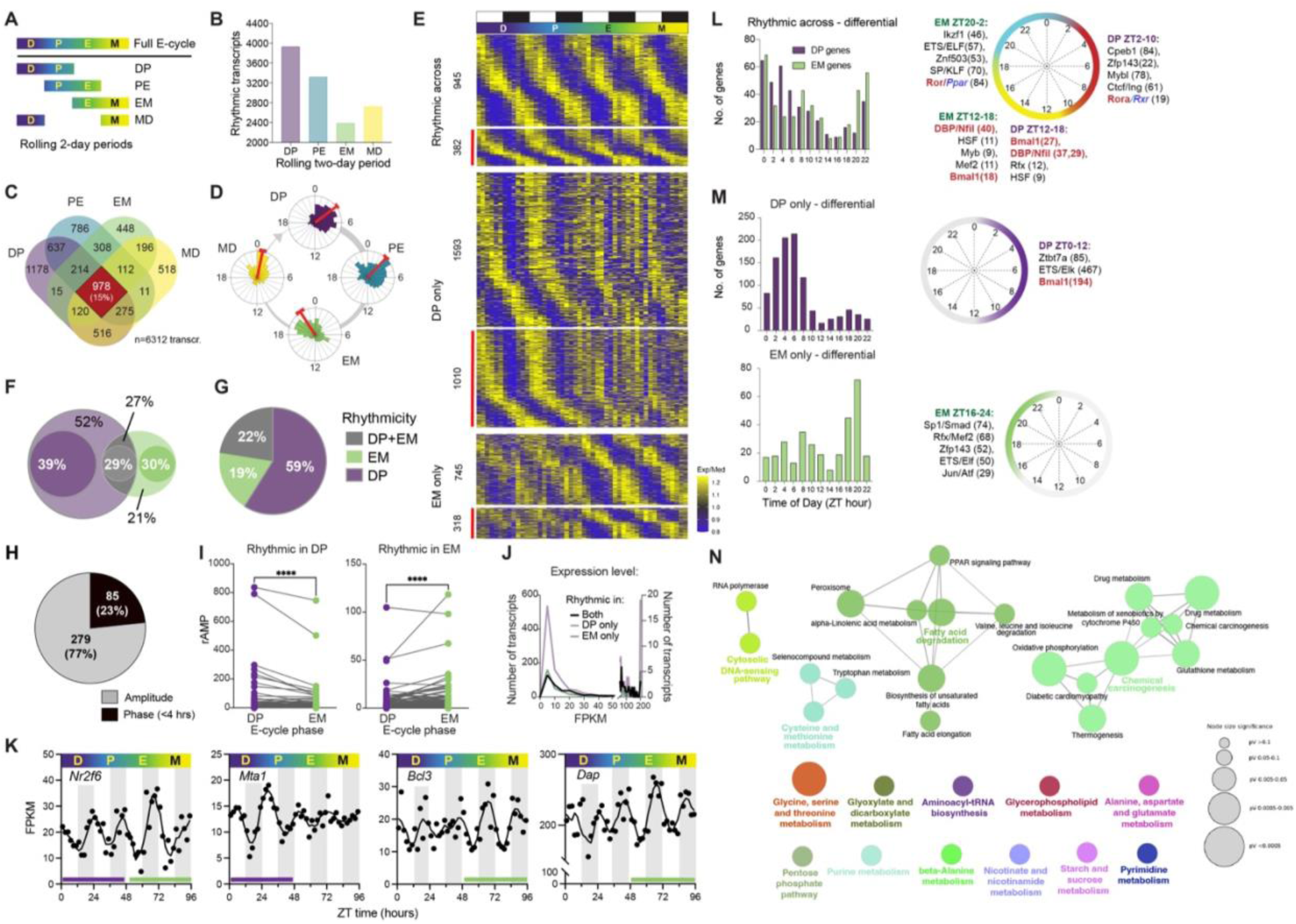
Estrous cycling impacts daily transcriptome cycling in liver. **A**. Rolling 2-day analysis paradigm. **B.** Numbers of rhythmic transcripts across each of the rolling 2-day periods. **C.** Venn diagram of the rhythmic genes between the two-day rolling periods. **D**. Raleigh plots of peak expression timing (phase, ZT hours) in each of the rolling 2-day intervals from B. The red bar indicates the average peak expression ZT time for each group. **E.** Heatmap of expression for genes rhythmic in across the estrous cycle, or in DP/EM only. All transcripts were rhythmic (JTK p<0.05) in at least one group, and differentially rhythmic transcripts identified by DoDR (meta.p <0.1) are indicated by the red line. The numbers of transcripts are indicated for each group. **F**. Venn diagram of rhythmic genes (JTK p<0.05) in DP and EM, with differential rhythmicity determined by DoDR (inset circles, meta.p <0.1, red lines in E). Percentages in black represent Venn proportions, those in white indicate proportion of each group displaying differential rhythmicity assessed by DoDR. Combined, DoDR indicates that 34.2% of rhythmic transcripts are differentially rhythmically expressed. **G.** Distribution of the number of differentially rhythmic transcripts that were rhythmic in DP and EM or both. **H.** The number and percent of genes rhythmic in both DP and EM but differential that have a phase difference greater than 4 hours. **I.** Change in amplitude of differentially rhythmic transcripts rhythmic in DP or EM (**** = p<0.0001, Wilcoxon matched-paired t-test). **J.** Frequency distribution of average daily expression levels for differentially rhythmic genes rhythmic in DP, EM or both groups. **K.** Example individual 4-day expression profiles of differentially rhythmic genes. Purple/green lines denote rhythmicity (at JTK p<0.05) in each period (DP or EM). **L.** Frequency distribution of genes according to peak timing/phase for genes that were rhythmic but differentially expressed (i.e. 382 near the top of E(red line) and the 29% in F) along with transcription factor binding site enrichment analysis of genes peaking at different time intervals across the day in both sexes. Representative transcription factors or classes for the top 5 enriched sites are shown, along with the numbers of gene targets. Red font highlights the enriched core circadian transcriptional regulators that are canonically responsible for driving peak expression at the indicated times of day, blue indicates nuclear hormone receptors. Identical analysis of non-differentially (“normally”) rhythmic genes is shown in Fig S3F. **M.** Data represented as in L but for differentially rhythmic genes rhythmic in either DP or EM. **N**. Pathway interaction networks of differentially rhythmic genes across DP and EM associated with liver metabolism (refer to Fig 2L-M). Bolded non-black text indicates top enriched pathways.

We first used qualitative, venn-based analyses to compare rhythmic gene populations across these rolling 48-hour intervals. At this level, we found that both the total numbers (Fig 2B) and the populations of rhythmic genes (Fig 2C) varied systematically across the e-cycle, in an apparent ∼4-day rhythm. This pattern persisted regardless of what threshold we used to define rhythmicity (Fig S3A-B). The largest population-level differences were between the non-overlapping DP and EM phases of the e-cycle (Fig 2B-D, Fig S3A-B) aligning with pre-ovulatory (DP) and post-ovulatory (EM) states. Strikingly, there was also a robust time-of-day dependency of the variation across e-cycle: DP appeared to promote rhythms that peak during the day, and EM promoted those that peak during the night (Fig 2D-E). Combined, these population-level analyses seemingly suggest that the circadian expression of a considerable proportion of rhythmic genes may be strongly influenced by estrous cycle, implying that circadian and e-cycles interact in some way in the liver to modulate daily gene expression patterns.

To gain better insights into these differences, we employed DoDR (21) to statistically compare rhythms across DP and EM at the individual transcript level. DoDR was developed to compare rhythms, enabling identification of those with significant differences in phase, amplitude and “noise” between two conditions. For our purposes, this will enable us to focus on the most robust “differentially rhythmic” transcripts for further analyses (Fig 2E-F, Fig S3E). This analysis revealed that ∼30-40% of transcripts rhythmic in either DP or EM, or both, displayed ‘differential’ rhythmicity (Fig 2F). Most of these were rhythmic only in DP, but there were differentially rhythmic transcripts among genes rhythmic only in EM as well as genes rhythmic in both DP and EM (Fig 2E-G, Dataset S1). Transcripts that were rhythmic only in DP or EM but were *not* statistically different by DoDR (see Fig 2E) may still have important biologically relevant differences in their rhythmicity, but we excluded them from further analyses due to this ambiguity. For comparison purposes, transcripts that were rhythmic in both e-cycle groups and not different by DoDR were considered non-differential, or ‘normally’ rhythmic.

Although some transcripts rhythmic in both DP and EM appeared to have >4-hour differences in phase, most of the differential rhythmicity between DP and EM appears to be due to a change in amplitude of individual rhythms (Fig 2H-I, see also Figs 2E,K). These differences were not unique to those with low overall abundances (Fig 2J), although a higher number of low-abundance transcripts appeared to oscillate in DP. Differentially expressed genes rhythmic only in DP or EM maintained the distinct day-time bias: many genes rhythmic in DP peaked during the day (ZT 0-12) (Fig 2E, M) and genes rhythmic in EM predominantly peaked at night but were more broadly distributed across the day compared to DP (Fig 2E, M). These time-of-day specific patterns were distinct from that of genes rhythmic in both DP and EM, regardless of if they were normally (Fig S3F) or differentially (Fig 2L) rhythmic across groups. These population-level differences and the time-of-day specificity are interesting as they suggest that the e-cycle may be interacting with discreet circadian transcriptional processes.

To explore these possible mechanisms, we explored the differentially rhythmic genes for enrichment of DNA response elements in their upstream regions (22). Here we focused on genes classified into four main groups as described above: the ‘normally’ rhythmic genes (Fig S3F), rhythmic in both DP and EM but differentially rhythmic by DoDR (Fig 2L), and those rhythmic only in either DP or EM and differentially rhythmic by DoDR (Fig 2M). The most conspicuous finding was that site for the central circadian transcription factors CLOCK/BMAL1 were highly enriched among daytime peaking genes rhythmic in DP only (Fig 2M). This day-time specificity correlates well with the time when CLOCK/BMAL1 is most transcriptionally active in the liver. These sites were also enriched among genes differential but rhythmic in both DP and EM (Fig 2L) but were not enriched among ‘normally’ rhythmic genes (Fig S3F). This alludes to the possibility that e-cycle might be modulating CLOCK/BMAL1 dependent rhythmic gene expression. However, expression of well-known CLOCK/BMAL1 targets within the clockwork itself (e.g., *Dbp*, *RevErbα*, Fig 1, see also Fig S4) was not detectably different between DP and EM. Taken together, these analyses predict that e-cycle may be modulating the rhythmicity in a target-gene specific manner, and not whole-scale modulation of the circadian clockwork activity itself.

Similarly, sites for the DBP/NFIL (drives early night peaking genes) and RevErb/ROR (drives late night peaking genes) sites were also enriched among both normal and differentially rhythmic genes (Fig S3F, Fig 2L). It should be noted however, that the ROR sites enriched in both the normal and differentially rhythmic genes share similarity with other nuclear hormone receptor (NHR) binding sites (as is the case with many NHRs; (23)), opening the door for potential direct involvement of estrogen or other hormones (note, *Nr3c3*, encoding nuclear progesterone receptor, is not detectably expressed in liver, see Dataset S1). RevErb/ROR sites were not, however, enriched in the population of nighttime peaking genes rhythmic in EM (Fig 2M). Instead, we found enrichment of sites for other transcription factors broad and diverse roles in this group (Fig 2M), many of which also enriched among genes peaking at other times of day or multiple groups (Fig 2L-M, including males – see Fig 3 and below). Thus, any time-of-day or e-cycle specific role for these other transcription factors is not clear. Nonetheless, *Dbp*, *Nfil3*, *RevErb,* and *Ror* genes appear normally expressed across the e-cycle (Fig1, Fig S4), implying that the e-cycle may modulate their function at specific downstream targets, independent of any potential regulation of CLOCK/BMAL1 discussed above. Combined, these findings suggest that e-cycles may modulate the activity of multiple core circadian transcriptional regulators in a target-gene specific manner. Future studies will be needed to determine if this is the case and decipher how these interactions operate.

**Figure 3.**
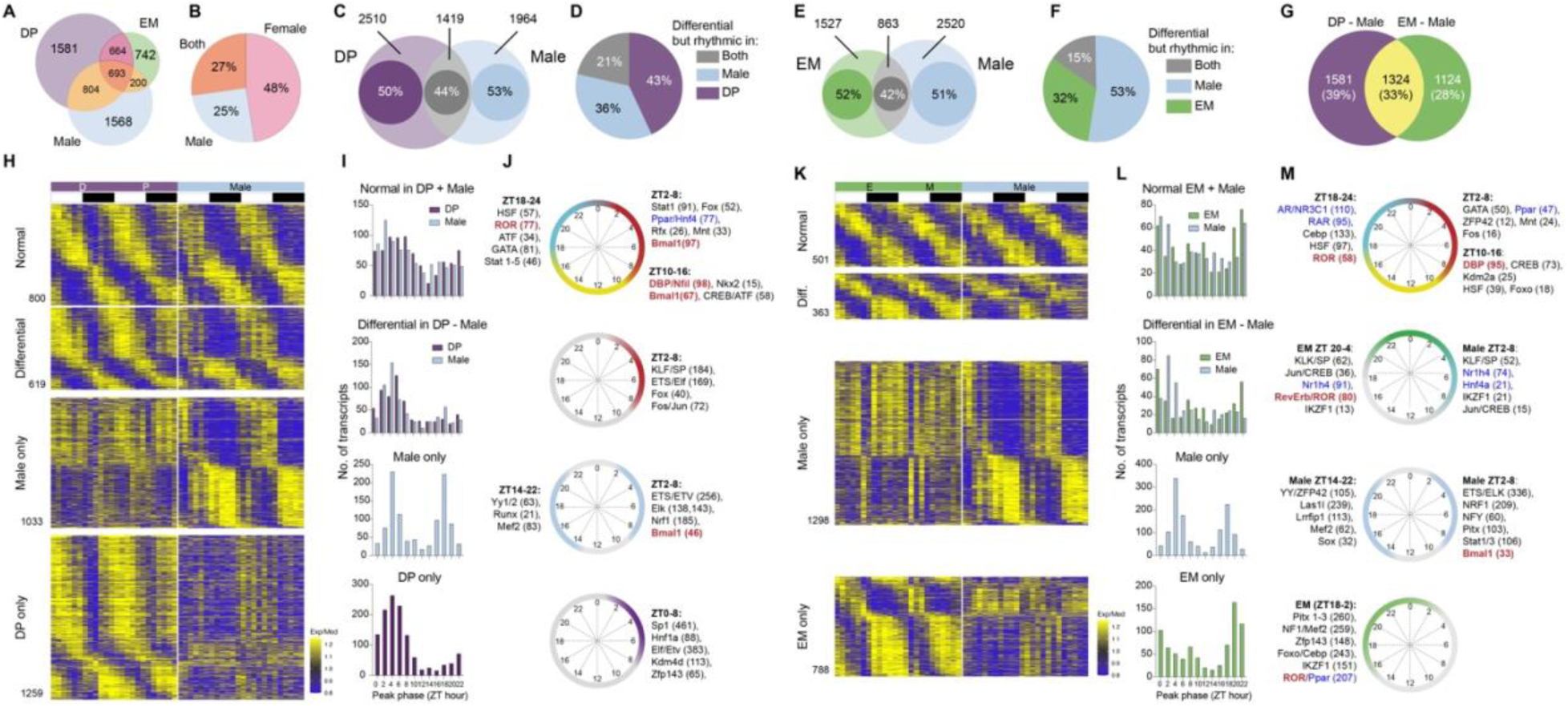
Estrous cycle expands sex differences in rhythmicity. **A.** Venn diagram comparing the population of rhythmic transcripts (JTK p<0.05) in males and the two rolling estrous cycle stages (DP and EM); the number of transcripts in each section is indicated. See also Fig S5. **B**. Pie chart showing the overall proportions of rhythmic transcripts detected that were male or female (DP and/or EM) specific. Here, if genes were rhythmic in DP or EM and males, they were counted as “both”. **C/E**. DM-Male (C) or EM-Male (E) comparisons using DoDR plotted as in Fig 2H. **D/F**. Proportions of differentially rhythmic genes (by DoDR) from C/E rhythmic in each group (JTK p<0.05). **G.** Venn comparison of all transcripts differentially rhythmic between males and DP and EM. Yellow represents genes rhythmic in both female groups and differentially rhythmic from males. **H/K.** Heatmap of expression of rhythmic genes from C/E, respectively. ‘Normal’ genes are those rhythmic in both groups but not different by DoDR (JTK p<0.05, meta.p >0.1). Data (FPKM) were median normalized across groups to enable comparison of relative expression levels. White lines are drawn on top to help separate groups. **I/L**. Frequency distributions of peak expression phase (ZT hours) for each comparison. **J/M**. Transcription factor binding site enrichment analysis, shown as described for Fig 2. Red font the enriched core circadian transcriptional regulators that are canonically responsible for driving peak expression at the indicated times of day. See also Figure S4-S5. Blue font highlights potential nuclear hormone receptor sites.

It is noteworthy that these differences strongly correlate to the days before (DP) and after (EM) ovulation, the key event controlled by the e-cycle, implying that the daily demands on liver function may fluctuate across the e-cycle. Indeed, differential rhythmicity across the e-cycle has the potential to influence metabolic pathways in the liver (Fig 2N). These phases correspond to substantially different sex-hormone environments (i.e. generally high estrogen, low progesterone in DP, reverse in EM), making them lead candidates in driving differential circadian regulation, either directly or indirectly. By extension, this suggests that disruption of circadian rhythmicity (e.g. shiftwork) may have differential consequences depending on the e-cycle phase or other conditions that impact the e-cycle (i.e. pregnancy, birth control, life-cycle stage), but this has yet to be explored.

### E-cycle broadens sex differences in rhythmicity

Previous work initially highlighted broad sex differences in the liver transcriptomes, as well as revealed some potential sex differences in the expression of clock genes [6,10–12,23]. This prompted us to compare how e-cycle impacts female gene expression compared to the rhythmic transcriptomes in males. We therefore collected males housed under identical conditions and paradigms to enable direct comparison to females in DP and EM (these data are in Dataset S2).

As anticipated, we observed substantial sex-dependent population-level differences in rhythmicity across the liver transcriptomes (Fig 3A-B, Fig S5). Comparable to what we and others have reported (6,10–12), we observed a > 1.5 fold increase in the population of rhythmic genes in female livers compared to males. Adding data from females also appears to at least double the total number of rhythmic genes overall in the mouse liver (Fig 3B, Fig S5B). Moreover, qualitative Venn-based analysis estimates that the rhythmic expression of as few as ∼27% of genes is conserved across sex (Fig 3B), with estrous cycle modulating this difference (Fig 3A).

As above, we employed DoDR to perform pairwise comparisons between DP and male (Fig 3C) or EM and male (Fig 3E) to more precisely identify differentially rhythmic genes across sex incorporating the e-cycle (Fig 3). Overall, ∼50% of genes rhythmic in *either* comparison (i.e. DP or male/EM or male) and >40% of genes rhythmic in *both* comparison groups (i.e. DP and male/EM and male) were identified as differentially rhythmic by DoDR (Fig 3C-D,E-F). Thus, this suggests that nearly half of the rhythmic transcriptome in liver is sex-dependent, regardless of e-cycle phase. Interestingly, the distributions of differentially rhythmic genes varied with e-cycle based comparisons (compare Fig 3D and F, and see below), mirroring the overall differential rhythmicity across e-cycle (Fig 2G). Approximately 33% of the differentially genes were common in both DP-male and EM-male comparisons (Fig 3G), potentially representing sex-chromosome based influences. The remaining ∼67% of differences appear to be modulated by e-cycle, presumably reflecting oscillating sex-hormone based influences. Combined, although the latter comparisons are indirect, these findings collectively reinforce that sex has a broad impact on rhythmicity in the liver and suggest that at least e-cycle hormones (and possibly others) may have distinct contributions to creating these differences.

We noticed that several core circadian clock genes were differentially rhythmic between males and females, with some apparently influenced by e-cycle phase (Fig S4). The most common difference was in overall expression level between the sexes, usually due to a difference in peak expression. These differences appear to follow distinct patterns specific to discreet loops and paralogs within the core clockwork (Fig S4). In the RRE loop, expression of RevErbα/β (repressors) appear higher in males, but expression of *Rorc* (activator) appears higher in females. In the D-box loop, both the activators (*Dbp*) and repressor (*Nfil3*) both appear more robustly rhythmic in females compared to males. In the E-box loop, the paralogs *Clock/Npas2* and *Dec1/Dec2* have one paralog that appears higher overall levels in males (*Clock, Dec2*), whereas the other paralog is higher in females (*Npas2, Dec1*). It should be noted that while we and others have observed several of these differences in previous studies (6,12), not all studies with male and female data have detected them (10,11). Also, it is not clear that these differences directly correlate to predictable differences in their overall function as their transcriptional targets within the circadian feedback loop system are not altered in predictable ways (Fig S4E). Nonetheless, the apparent patterns to these differences suggest they may be intentional and underscore the need for a deeper understanding of overall sex-differences in rhythmicity throughout the organism.

Beyond the clockwork, prominent differences were seen in genes that were rhythmic in only one of each of the comparison pairs (DP – Male, Fig 3H-I or EM – Male, Fig 3K-L). Genes rhythmic in only males were expressed nearly exclusively in two primary clusters – one with peak expression around midday, the other peaking around midnight. The same genes in females have severely damped rhythms by comparison, but their overall relative expression, high or low, in females strongly correlated with the time of day they peaked in males (Fig 3H,K, see also Fig S6E-F). Daytime peaking genes in males were constitutively expressed at near their peak levels in both DP and EM, but nighttime peaking genes in males were expressed at near trough levels in both female groups. Since the pattern appears similar in both DP and EM comparisons, we predict that sex is likely a major factor in this difference.

Similarly, genes rhythmic in DP or EM, but not males, also had prominent clustering around mid-day and mid-night, but these clusters were e-cycle phase specific: the daytime cluster was present in DP (Fig 3H-I bottom), and the nighttime cluster was specific EM (Fig 3K-L, bottom). In contrast to the pattern described above, expression of these genes in males was relatively low overall, regardless of when they peaked in females (bottom panels of Fig 3I, L). While these were the most prominent patterns, other potential patterns are apparent (Fig 3H-I and K-L), likely owing to a broader distribution of rhythmic genes across the day in females than males, particularly in EM (see also Fig S5). Thus, it appears that the mechanisms promoting rhythmicity in females may be more complex than males in this context. Nonetheless, the strong daytime component in DP and strong nighttime component in EM suggest that the e-cycle may be modulating as few as two different rhythmic transcriptional activation mechanisms in females that are non-functional in males.

Rhythmic expression of genes encoding transcription factors appears to be broadly modulated by sex and e-cycle (Fig S7). Therefore, we again identified potential candidate transcriptional pathways that may be involved in producing the different transcriptional patterns, as above (Fig 3J,M). Genes that were normally rhythmically expressed in males and DP or EM were enriched for the core circadian clockwork transcription factors as expected (Fig 3J top, Fig 3M top). The only exception to this was that we did not find enrichment for CLOCK:BMAL1 sites in genes rhythmic in both males and EM; however this may be due to differences in the time-of-day distribution of rhythmic genes in this comparison (i.e. low number of day-time peaking genes in EM; Fig 3L, Fig S6).

Among the differentially rhythmic genes, we found enrichment for sites bound by several factors, but most were not specific to a particular sex/group or time of day (black font in Fig 3J,M). We did find some enrichment of CLOCK:BMAL1 sites in daytime peaking genes rhythmic only in males compared to DP or EM (Fig 3J,M, third panels down). Oddly, these sites were not enriched among the daytime peaking genes in DP (Fig 3G bottom) as they were in DP – EM comparisons (Fig 2), but they were strongly enriched in this group defined by rhythmicity alone (without using DoDR; Fig S5H). In addition, nighttime peaking genes differentially rhythmic between males and EM were enriched for sites for RevErb/ROR and other nuclear hormone receptors (Fig 3M) reminiscent of the e-cycle differences (Fig 2). Overall, these analyses reveal that while there are strikingly distinct differential expression patterns, and the potential for involvement of some circadian clockwork factors at some target genes, candidate transcription factors that may drive the bulk of these differences remain unidentified.

### Downstream health implications for estrous cycling and the clock

It is clear, however, that there are marked influences of e-cycle on sex-dependent circadian rhythmicity in the liver transcriptome (Fig 3). To explore the potential consequences of these differences, we assessed how these variations might translate physiologically in the liver. Overall, roughly 47% of all liver-specific metabolism and disease related genes (25) are differentially rhythmic in our combined datasets, where differential rhythmicity specific to e-cycle phase appears to have a major impact (Fig 4A, see also Fig S8). Non-alcoholic fatty liver disease (NAFLD) is a prime example as it represents both liver metabolism and disease, and is sensitive to circadian dysfunction (i.e. shift-work, late-night eating). Our data suggests that rhythmic expression of up to ∼75% of NAFLD-related genes (26,27) may be influenced by sex and/or e-cycle (Fig 4B-C, Fig S8B), implying that circadian-related NAFLD development may occur via different mechanisms in males and females, and this might vary across the e-cycle. Thus, our data reveal that e-cycle modulation of rhythmicity in females, in addition to sex, adds new dimensions to potential mechanisms and pathways leading to liver disease progression and treatment. Given that circadian rhythms are pervasive throughout the body, we predict that these findings from the liver are representative of the potential differential circadian regulation across organisms and the overall disease and therapeutic landscape.

**Figure 4.**
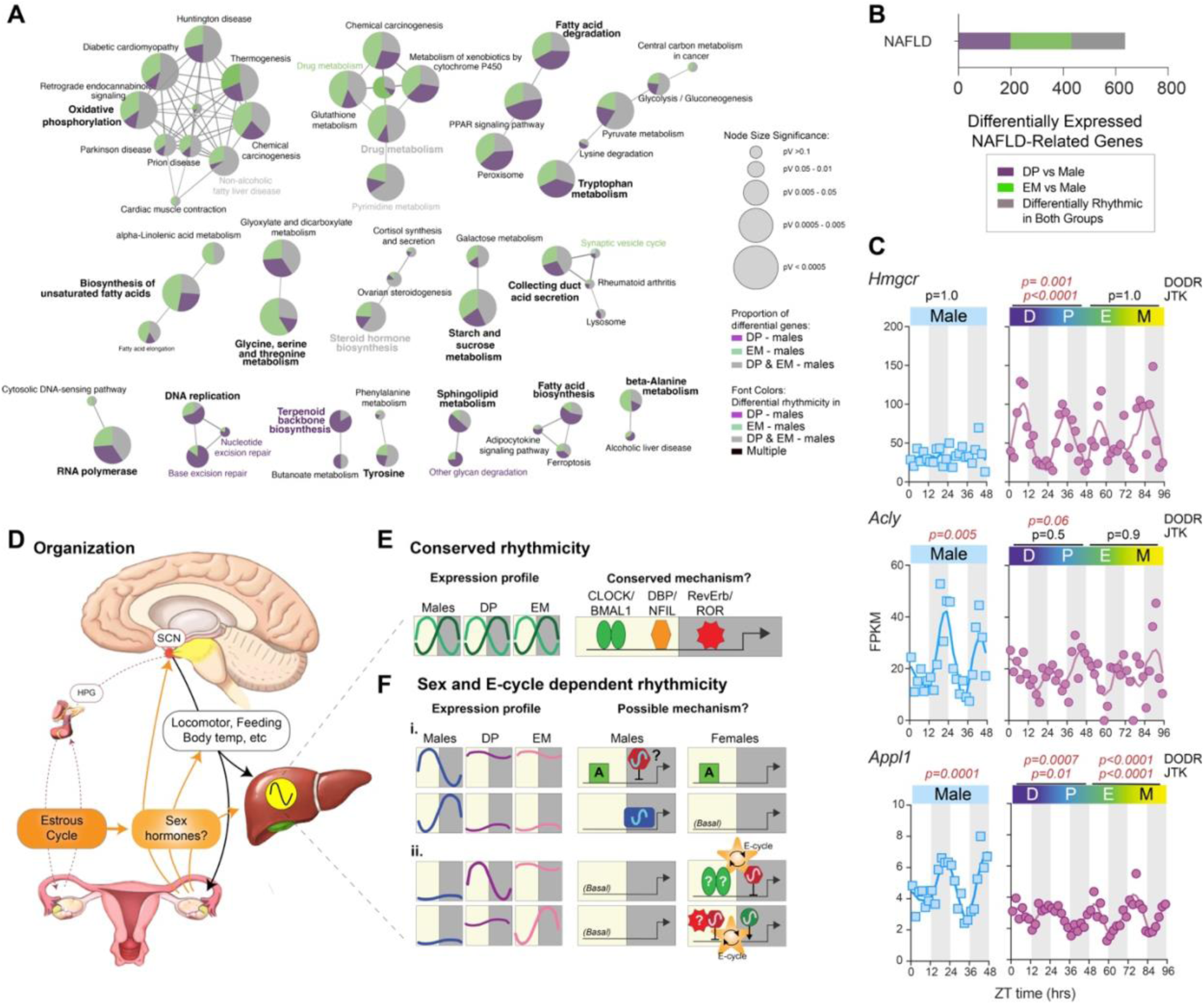
Estrous cycle plays a role in sex differences with metabolic interactions. **A.** Pathway interaction networks (as in Fig 2) of differentially rhythmic genes depicted in Fig 3. Bolded text indicates top enriched pathways. **B.** Distribution bar plot of differentially rhythmic genes associated with NAFLD (45,46). **C.** Example individual expression profiles for genes associated with NAFLD. Significant DoDR meta.p (to males) and JTK p-values are indicated above (red font = <0.1 or 0.05, respectively). **D**. Schematic of organization and intersecting points (arrows) between e-cycle and circadian regulation across the body (tissue cartoons were created using ChatGPT). **E.** Cartoon representation of rhythmic gene expression profiles that appear conserved across sex and e-cycle (left) and the potential core circadian loop elements driving these rhythms (day = yellow/night = gray shading). **Fi.** Model for male-specific rhythmicity. Daytime = strong constitutive activation (green box labeled A) in both sexes, rhythmicity in male driven by rhythmic repression at night (red hexagon). Nighttime = male specific nighttime activation (blue box). **Fii.** Model for female-specific rhythmicity. Female/DP daytime = Low basal activation in males but strong activation (green ovals, possibly CLOCK:BMAL1) in females, with rhythmic repression (red hexagon) that is regulated by e-cycle (orange star). Female/EM nighttime = unknown female-specific rhythmic nighttime activators (green circle) or daytime repressors (red hexagon, possibly REV/ROR) that are regulated by estrous cycle (orange star).

## Discussion

There is a long history of evidence that circadian rhythms can be influenced by sex in mammals. The consensus hypothesis has been that the circadian system largely fulfills the same overall functions in both sexes. These aspects relate to controlling sleep-wake and metabolic and other physiological rhythms, and the main sex differences are centered around reproductive biology. However, with the detection of sex-specific transcriptional mechanisms closely associated with the clock in the liver (Fig S4) (6,10–12), it is becoming increasingly evident that the circadian system has evolved strikingly different roles in males and females. Moreover, many studies are performed in reproductive-age adult animals, but the complex differential hormonal environments between the sexes are usually not accounted for because to do so would challenge experimental feasibility. One of the goals of this study was to see what could be missing, if anything, by not accounting for e-cycles at the transcriptional level. Much to our surprise (and some chagrin), the data presented here suggest that we have been indeed, missing quite a bit, as only ∼36% of rhythmic genes in females appear ‘normal’ across, with the differential populations showing distinctly time-of-day specificities (Fig 2).

This presumably occurs via interactions between circadian and estrous cycles as they interact at many points (Fig 4D) (9). The central circadian clockwork, via the HPG, influences most female sex hormones, and gates ovulation during the night in mice. Moreover, photoperiodic/seasonal breeding patterns require a functioning circadian clock system (28), and ovulation itself requires a functioning molecular circadian clockwork in the ovary (29–31). In turn, circadian locomotor, feeding, and body temperature patterns are also modulated by e-cycle hormones, creating potential feedback loops between the two cycles at multiple points (Fig 4D)(4,5,32). Each of these physiological rhythms can also regulate circadian gene expression in the liver, thus it is possible that they all may contribute to differential rhythmicity.

What is not clear, at least from our data, is if the ∼4-day cycles in sex-hormones themselves directly contribute to circadian gene expression in the liver. GPCR-based receptors for estrogen, LH, FSH, and progesterone are expressed in the liver (Dataset S1), thus each could possibly signal an e-cycle phase to the liver. Indeed, binding sites for transcription factor effectors of cell-signaling cascades were enriched across genes with differential rhythmicity profiles (Figs 2-3), but these factors are also likely integrated with other cell-signaling events, which we predict would likely blunt the input of hormone actions at their GPCR receptor types. Nuclear sex-hormone receptors in contrast could have a direct role modulating rhythmic gene expression. Indeed, nuclear hormone receptor binding sites were enriched among differentially rhythmic genes, suggesting potential roles, although which receptors, or in which capacity, is not yet clear. Interestingly, nuclear progesterone and estrogen receptors can cooperate to coordinate e-cycle dependent changes in chromatin architecture that directly impact the activity of other transcription factors to cause differential gene expression in the uterus (33). Whether a similar mechanism could occur in the liver is unknown, but nuclear progesterone receptors are not detectably expressed in female mouse liver (Dataset S1). However, given that the circadian transcriptome appears modulated by e-cycle, it seems likely that a similar type of mechanism involving estrogen receptors ESR1/2 could be involved.

On the other hand, the highly consolidated differences support the robustness of the interaction and suggest that the transcriptional mechanisms underlying these differences may be quite discreet across time of day (Fig 4E-F). In some contexts, there are clear patterns that suggest candidate transcriptional mechanisms. For example, the difference between genes that peak during the day in males and are arrhythmic in females may be as simple as the presence/absence (respectively) of a nighttime transcriptional repressor (Fig 4Fi). Likewise, the rhythmicity of nighttime peaking genes in males could be driven by an activator that is only active at those genes in males.

The situation in females may involve two mechanisms (Fig 4Fii), one that influences rhythmicity by sex, the other by e-cycle stage. Based solely on relative expression levels, it appears that genes rhythmic in females are lacking robust transcriptional activation in males, fitting a model in which the circadian enhancer sites are only accessible/active in females. Indeed, the liver has sexually dimorphic DNA methylation and histone acetylation/methylation patterns that can lead to altered chromatin structures and functionally impact gene expression (6,34). Subsequent e-cycle regulation may target circadian regulation, by modulating the efficacy of nighttime repression for daytime-peaking genes in DP or activation/repression for nighttime genes in EM (Fig 4Fii). For these genes, at least in DP, our analyses suggest that sex and e-cycle may be modulating how the core CLOCK:BMAL1 factors either interact with or regulate specific target genes in females.

Once these mechanisms start to become identified, it will be interesting to determine how they are established in the first place. For example, it will be informative to determine if these differences are the result of developmental adaptations driven by the sexual maturation process (i.e. possibly epigenomic?), rather than active/direct modulation by sex-hormone receptors or sex-chromosome linked factors (6). In addition, it will be important to determine if these change with female life-cycle and/or reproductive status (i.e. pregnancy or hormone-based birth control) – if they indeed are driven by sex hormones, then differential rhythmicity could be strikingly different across the female lifespan. Determining the underlying drivers and mechanisms will undoubtedly provide key insights into female circadian biology, as well as assisting in designing precise experiments to assess their impacts, as incorporating the full e-cycles in all experiments is not always feasible.

Finally, our data also have direct implications for liver disease and therapeutics among other functions. It is becoming well established that circadian systems in females are more robust and resilient to perturbation compared to males, at least some of which is linked to estrogen signaling and the gut microbiome (6,9,35,36). Our findings also suggest that this robustness might vary with e-cycle phase, as at least in the liver, rhythmicity is generally less robust and is shifted to a nighttime bias in the phases where estrogen is low (i.e. EM). In addition to providing insight to the intersection between time-of-day and sex, these data also provide advancement for tailoring therapeutic interventions to individuals’ endogenous conditions. These data (Datasets S1-S2) serve as an initial resource enabling groups to start evaluating if e-cycle is a factor for consideration in their studies, in addition to providing starting points for identifying novel interactions between e-cycle and liver-related physiology. Ultimately, however, we predict these findings will extend beyond the liver as nearly all tissues display robust circadian rhythmicity.

## Materials and Methods

### Animals and Diets

Adult female and male C57BL6 mice were obtained from Jackson Laboratory (stock number 000664) at 9-10 weeks of age. This strain was used due to its wide availability and usage. After a 3-day acclimation period, they were individually housed under a 12:12 light/dark cycle, with *ad libitum* access to water and food (PicoLab Laboratory Rodent Diet 5L0D, LabDiet, St. Louis MO, USA), and allowed to further acclimate to for an additional week before either e-cycle staging (females) or tissue collection (males). At the indicated times, mice were euthanized using CO2 followed by decapitation to collect trunk blood and dissect tissues. All animal studies were approved by the Institutional Animal Care and Use Committee of Morehouse School of Medicine in accordance with the United States Public Health Service Policy on Humane Care and Use of Laboratory Animals (protocol numbers 21–14, 24-10).

### Estrous Cycle Staging and Animal Collection

Estrus cycle stage was identified initially using cytology of vaginal lavages (37). Briefly, each female mouse was vaginally lavaged with 1x PBS, and cytology was imaged using an inverted light microscope and used to assign a preliminary e-cycle phase. Each animal was lavaged daily, at the same time each day up to the time of serum and tissue collection. Animals were maintained on the LD cycle throughout to eliminate any potential drift caused by differences in individual free running or any unknown resetting effects of the daily vaginal lavages.

Each animal’s serum and tissues were collected at 2-hour intervals across each day, from animals that were in each of the four e-cycle stages (diestrus, proestrus, estrus, metestrus), in a discontinuous fashion. Most of the 70 mice we were collected during the first 2 days, but sampling became more sporadic as we waited for mice to enter the right e-cycle phase; All animals/tissues were collected within a 7-day period. Male mice were collected continuously every 2 hours over a 2-day period. Isolated serum (from trunk blood) and dissected livers were snap frozen in dry-ice and stored at -80C until processing.

### RNA Isolation and Sequencing

Liver samples were processed for RNAseq as we have done previously (12). Briefly, total RNA was extracted from the liver using Trizol reagent (Invitrogen) per the manufacturer’s instructions, followed by an RNA clean up using Rneasy kits (Qiagen). RNA (500 ng/ul) was used to generate strand-specific, Poly-A + RNAseq libraries. RNAs were converted into sequencing libraries by using Illumina TruSeq stranded mRNA Library Prep kits and sequenced by Omega BioServices (Norcross, GA) using the Illumina HiSeqX10 platform. Samples were sequenced to a depth >35 million 150bp X 150bp paired end reads. The reads were mapped to the mouse MGSCv39-mm9 genome using Tophat 2.1.0 and fragments counted and calculated using Cufflinks 2.2.1. Low expression genes were filtered out if the sum of their raw count was less than 100 across the time points in all groups (female and male). Count data was used for DESeq2 (https://yanli.shinyapps.io/DEApp/), and FPKM was used for all other analyses. The female sample ZT 25 (proestrus ZT1) gave RNAseq data that were substantially out of line with neighboring timepoints, which created a major artefact in circadian analyses, despite resequencing and mapping etc; it was therefore eliminated from circadian analyses and replaced with the average of the point before/after in all plots for continuity. All Raw (FASTQ) and processed data (counts, FPKM= fragments per kilobase per million) are available at NCBI BioProject (PRJNA1348461) and the processed FPKM data are available in Supplemental Datasets S1-S2.

### Serum hormone analyses

Sex hormones were measured from serum collected, and stored at -80C, from harvested females and submitted to Ligand Assay & Analysis Core at the University of Virginia (https://med.virginia.edu/research-in-reproduction/ligand-assay-analysis-core) as we have done previously (12). In some instances, we harvested up to 3 mice at each time/e-cycle stage and ultimately selected the individual with serum hormone profile that best matched expectations for e-cycle. In the few instances where the hormone data was inconsistent with the vaginal cytology, the animal was either excluded from the study (i.e. not subjected to RNAseq), or the e-cycle stage was reassigned based on the hormone data.

Correlations between transcript FPKM and serum FSH or estradiol levels were performed on median-normalized levels. We determined Pearson correlation coefficients (r) for each of the transcripts to each hormone across the 47 samples (ZT25 was excluded). Correlations were performed utilizing a custom script written in the Igor Pro 9 (Wavemetrics, Lake Oswego, Oregon) which calls the Igor Pro function *statsLinearCorrelationTest* which determines the Pearson Correlation Coefficient for two sets of data. The minimum r value that reached statistical significance (p<0.05) was 0.288; however, this was not corrected for >20,000 multiple tests. Therefore, we focused on transcripts with more robust correlations, using r =0.4/-0.4 as a cutoff. The full list of r values for all genes with each hormone are in Dataset S1.

### Time Series Analysis for Circadian Profiling

We used Nitecap to assess circadian expression profiles, using JTK outputs to provide circadian measures (p/q-values for rhythmicity, peak phase, amplitude) (https://nitecap.org) (15,38). As noted above, JTK cycle is robust against outliers, and appears generally more conservative at identifying rhythmic transcripts compared to other methods, as it can be sensitive to ‘noise’ in the data (20). In most figures in the main manuscript, rhythmic transcripts in each group were identified using a relatively low stringency JTK p-value threshold of 0.05 to help reduce overestimating differences between groups, while still considering less robustly rhythmic transcripts (15), however increasing thresholds and controlling for false-discoveries yielded similar proportional results (Fig S3, S5), just with reduced populations of rhythmic genes. We employed qualitative Venn-based analysis as an initial overview comparison of the transcriptome groups, as defining ‘rhythmicity’ is an essential component for downstream analysis by DoDR (21). We chose DoDR for these comparisons because it employs multiple algorithms to identify differences, and its overall results agree with most other algorithms developed for similar purposes (39). DoDR analysis was performed using transcripts that were defined as rhythmic (JTK p<0.05 or q<0.05-see Fig S3E) in either or both comparison groups, and were considered ‘differentially rhythmic’ if they had a DoDR meta.p value less than 0.1(39). Genes were categorized based on their rhythmicity in one or both groups and difference detected by DoDR (JTK + DoDR). Overall, we provide all the data in Supplemental Datasets S1-S2 and encourage others to explore these data as a resource for potential future studies.

Oriana (version 3, Kovach Computing, Anglesey, Wales UK) was used to produce Raleigh plots. GraphPad Prism (v10.4.0; GraphPad Software, SanDiego CA, USA) was used for further analysis and plotting. Heat maps were generated using an R code: (https://github.com/gangwug/SRBR_SMTSAworkshop/blob/master/R/fig.R). Gene lists were imported into gProfiler for gene ontogeny/pathway analysis (https://biit.cs.ut.ee/gprofiler/gost) (40,41) and KEGG where applicable (https://www.kegg.jp/kegg/pathway.html) (42). Venn diagrams were created using the webtool at https://bioinformatics.psb.ugent.be/webtools/Venn/. Gene lists for liver transcription factors were identified from https://cgs.csail.mit.edu/ReprogrammingRecovery/mouse_tf_list.html (43).

### Transcription factor binding site enrichment analysis

Promoter enrichment analysis was performed using iRegulon (version 1.3) (22) as an app within Cytoscape (version 3.10.2) using the default parameters: 20kb upstream of the transcription start site; 10k (9713 PWMs) motif collection; 0.03 ROC threshold for area under the curve; 0.0001 FDR. Networks of candidate transcription factors and their target genes were generated in Cytoscape. ChIP predicted targets were used to compare candidate BMAL1, CLOCK, and NPAS2 target genes from iRegulon (see Fig 3, S10) (44).

### Statistical analysis

All statistically analyses including those for circadian rhythmic detection and promoter enrichment were performed using the embedded analysis in each algorithm (see figure legends) or using GraphPad Prism (v10.4.0; GraphPad Software, SanDiego CA, USA) if not otherwise stated.

### Metabolic Interaction Analysis

The dataset for liver metabolism related genes was obtained from the Knepper Lab of the NHLBI (https://esbl.nhlbi.nih.gov/Databases/KSBP2/Targets/Lists/MetabolicEnzymes/MetabolicEnzymeDatabase.html) (25). The NAFLD related datasets were obtained from the Maayan Lab https://maayanlab.cloud/Harmonizome/gene_set/Non-alcoholic+Fatty+Liver+Disease/CTD+Gene-Disease+Associations) (45,46). The plug-ins, ClueGO (version 2.5.10) and CluePedia(version 1.5.10) as apps within Cytoscape (version 3.10.2), were used for KEGG analysis to indicate top enriched pathways.

## Data Availability

All raw and processed data are available at NCBI BioProject (PRJNA1348461). The processed FPKM data are available in Datasets S1-S2.

## Acknowledgments

This work was supported by the NIH NIGMS grant R35GM127044 (JPD) and NIH NIMHD RCMI Grant U54MD007602 (Morehouse School of Medicine). HAD is also supported by NIH NIGMS Grant R16GM146703, and MB is supported in part by Simons Foundation grant SFI-AN-ECI-00009801 and NIH NINDS Grant R25NS117365. JPD, HAD, and MB are also supported by NSF EES CREST Center Grant 2514598. KAS was supported by HHMI Gilliam Fellowship GT16514. The content is solely the responsibility of the authors and does not necessarily represent the official views of the NIH, NSF, Simons Foundation or HHMI.

