## Supplemental Figures for "Remodeling of the hepatic circadian transcriptome across the estrous cycle"

**Supplementary Figures: Smith et al.**

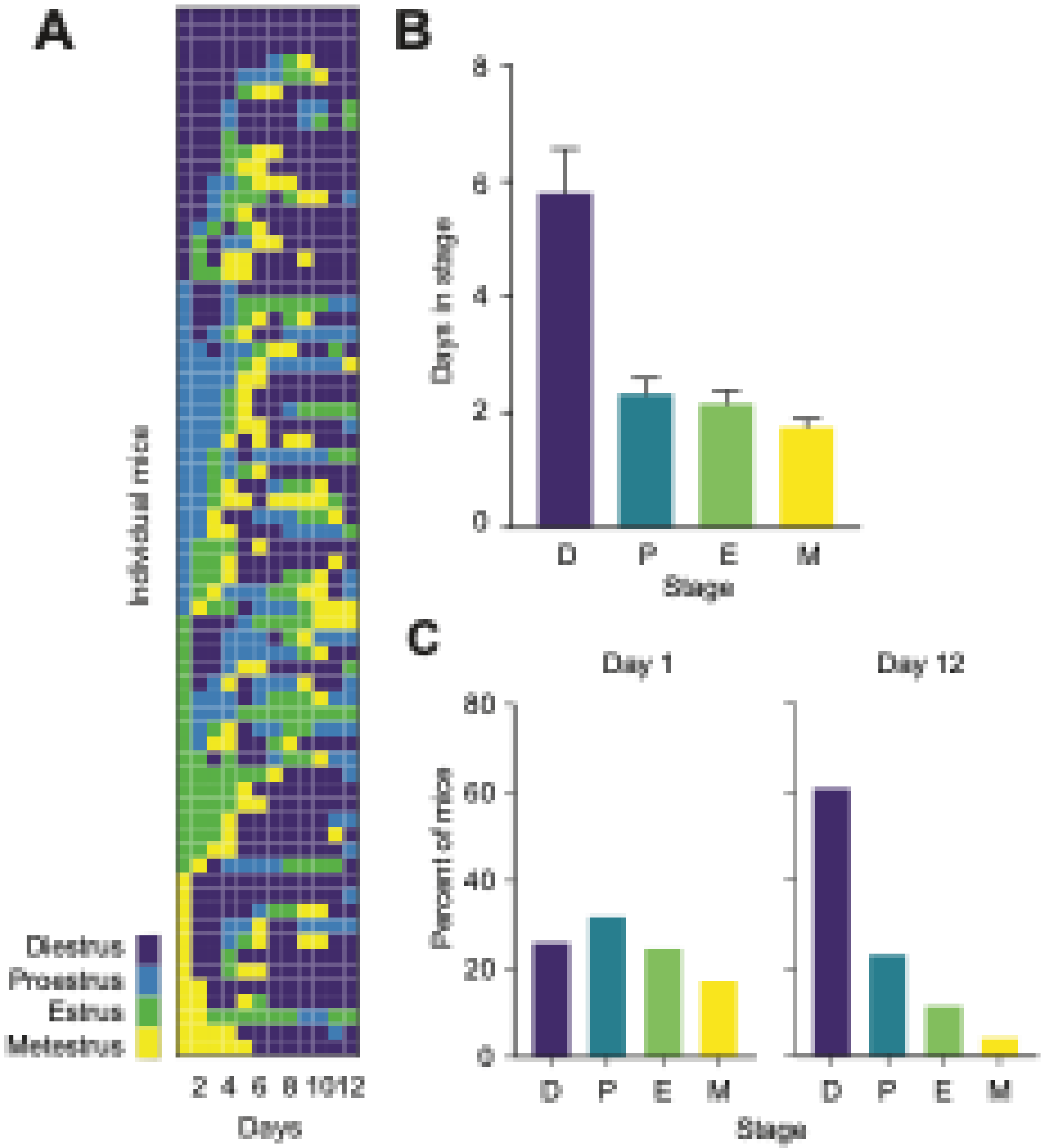


**Figure S1. Estrous cycling in individual C57Bl6 mice tracked daily for 12 days. A**. Heat map representation of estrous cycling in each of 69 individually housed C57BL6 mice over 12 days. Staging was determined from cytology of vaginal smears taken daily from ~ZT 2-4. **B**. Average (+/- 95%CI) number of days each mouse spent in each estrus cycle phase during the 12-day recording period. **C**. Percent of mice in each estrous cycle phase on the 1st and 12th day of monitoring.

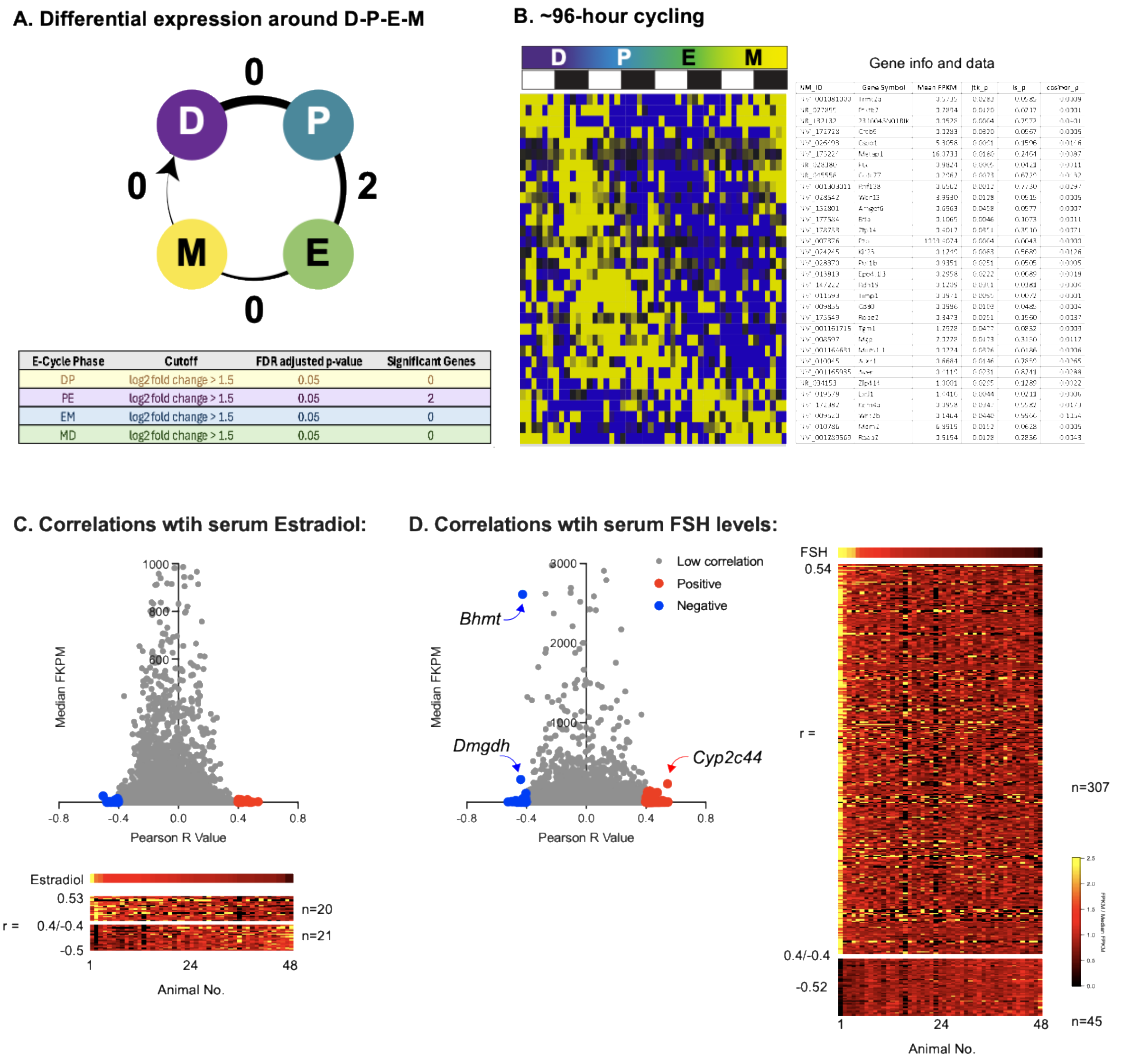


**Figure S2. Some genes exhibit cycling coincident with estrous cycle. A.** Differential (Deseq2) expression around D-P-E-M. All samples from collapsed across time for each e-cycle phase and compared in a pairwise manner (D vs P, P vs E, etc). Table below summarizes results. **B.** Heatmap of the 32 transcripts that display expression that follows e-cycle, determined using Nitecap (see Methods) tuned to identify transcripts with a ~96-hour period. Heatmap is organized by peak phase, and expression levels are median normalized to enable comparison of rhythm across genes. Individual transcript IDs are listed to the right (lined up with the heatmap), and lists NM ID, Gene ID, mean 96-hr expression (FPKM) and p-values form three rhythm algorithms (Nitecap). **C.** Volcano plots of FPKM and r values, red/blue = liver transcripts that were correlated (r >|0.4|) with serum estradiol (top); heatmap of relative hormone and mRNA levels for those with r >|0.4| (bottom). **D.** Genes whose expression was correlated with serum FSH levels, plotted as in C.

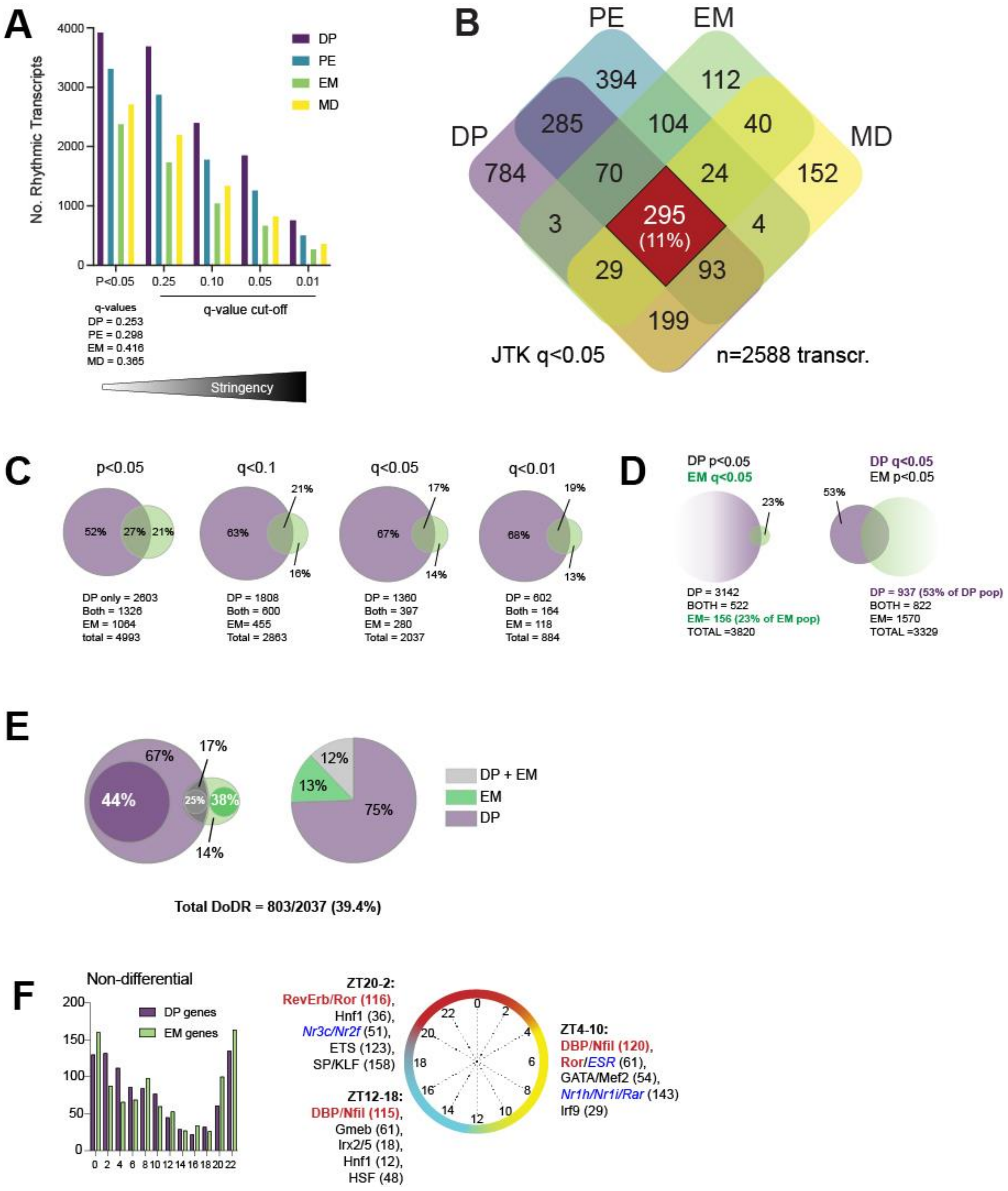


**Figure S3. Differential circadian rhythmicity suggests the clock may have different impacts across the e-cycle and persists across a range of analysis parameters. A**. Comparison of the number of "rhythmic" transcripts at different JTK cutoff values. The p <0.05 are the same as in Fig 2, and the corresponding q-values (Benjamini-Hochman false-discovery rate) at p=0.05 are shown for each group below.

Numbers of rhythm transcripts at increasingly stringent q-values are also shown. All values are derived from JTK analyses. **B**. Four-way Venn diagram (as in Fig 2C) of rhythmic transcripts using JTK $q<0.05$ as a rhythm threshold. **C.** Venn-based comparisons of the rhythmic gene populations between DP and EM at different rhythmicity cutoff levels (from B). Actual gene numbers are listed below; percent of total population is indicated within the diagram. **D.** Comparison of DP and EM using low ($p<0.05$) and high ($q <0.05$) thresholds shows that many genes meeting the high rhythmicity thresholds were still not rhythmic in other group at low thresholds (indicated by bold green or purple font). **E**. DoDR evaluation of differential rhythmicity between DP and EM similar to Fig 2 but using high ($q<0.05$) rhythmicity thresholds. Venn diagrams (replotted from C) are overlaid with darker circles depicting the proportions with a DoDR meta.p <0.1. **F**. Left: Frequency distributions of peak expression phase (ZT hours) for non-differentially (i.e. 'normally') rhythmic genes across e-cycle (DP and EM). Right: Transcription factor binding site enrichment analysis (iRegulon) of genes peaking at different time intervals across the day in DP and EM. The top 5 enriched sites (defined by a representative transcription factor or class; numbers of gene targets) are shown in brackets. Red highlights the enriched core circadian transcriptional regulators that are canonically responsible for driving peak expression at the indicated times of day. Blue font highlights sites for indicated nuclear hormone receptors.

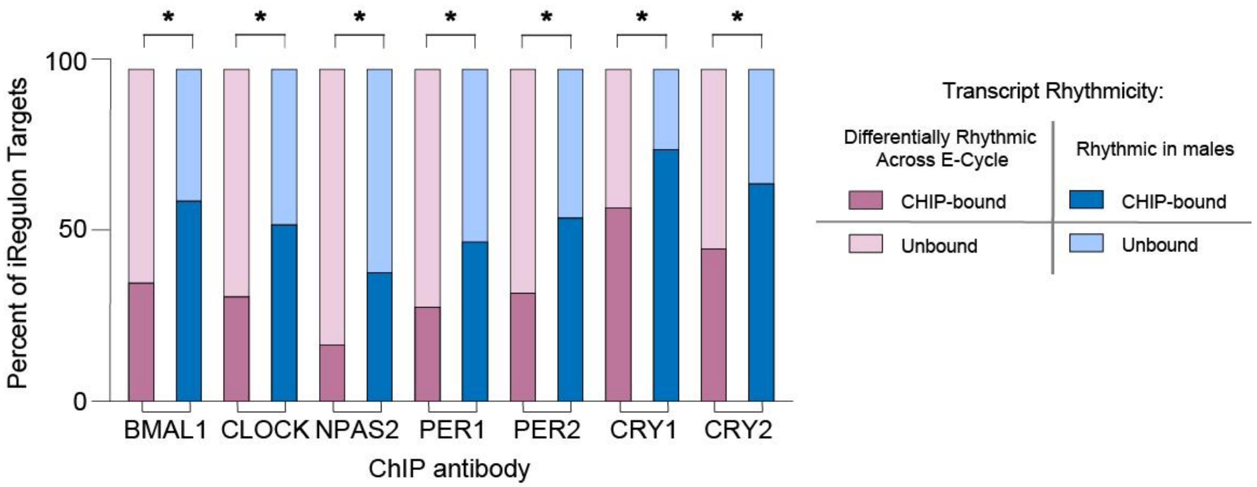


**Figure S4. Comparison of predicted DNA binding and published male circadian cistrome. A.** Bar plot of the percent of E-box gene targets bound to BMAL1, CLOCK, NPAS2, PER1/2, and CRY1/2 in males [1] Pink= differentially rhythmic across e-cycles; Blue= rhythmic in males (See Fig 2 for e-cycle data and Fig 3 male data).

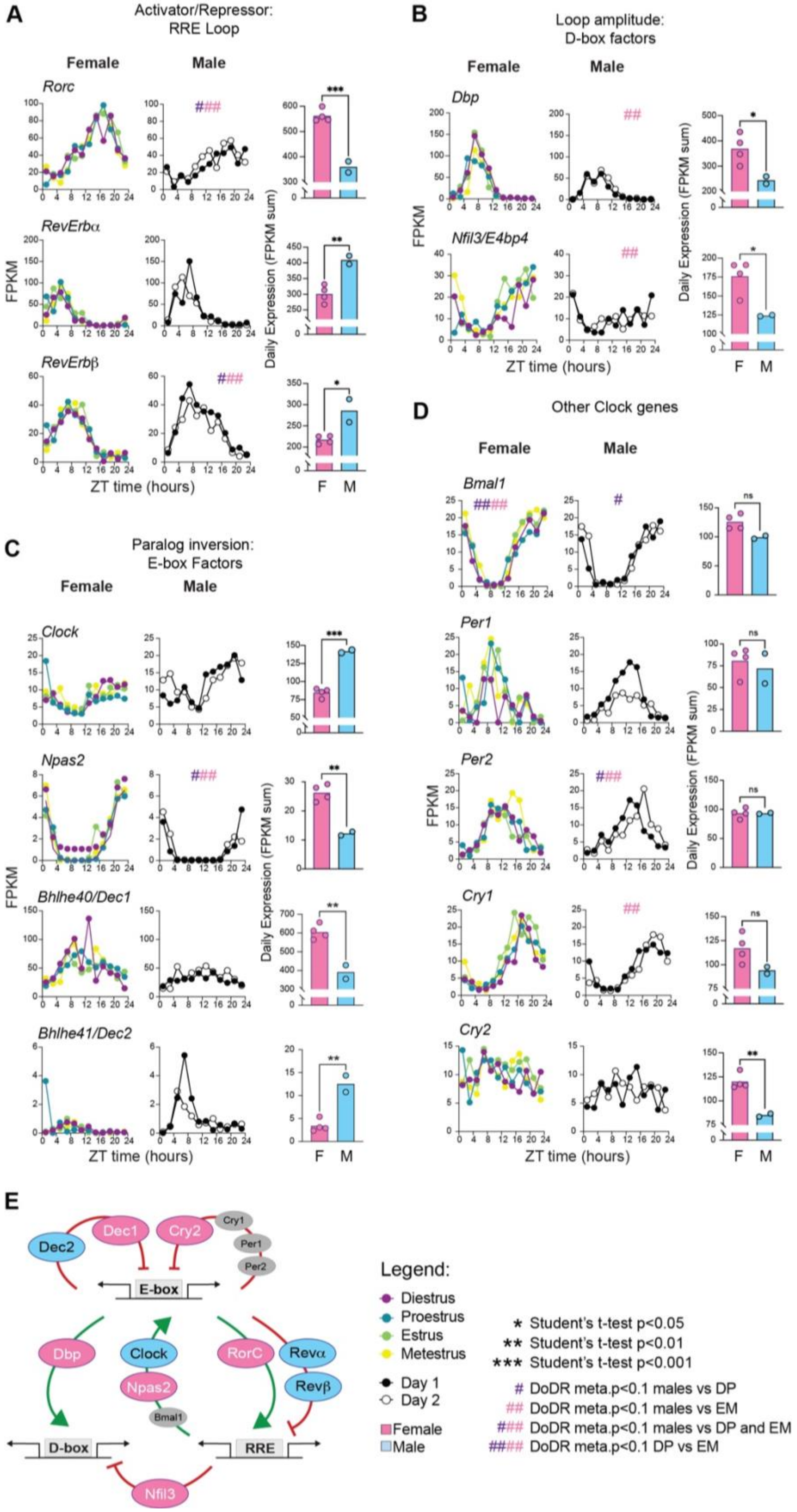
A
Activator/Repressor:
RRE Loop
Female
Male
Rorc
RevErbα
RevErbβ
FPKM
ZT time (hours)
Daily Expression (FPKM sum)
F
M
B
Loop amplitude:
D-box factors
Female
Male
Dbp
Nfil3/E4bp4
FPKM
ZT time (hours)
Daily Expression (FPKM sum)
F
M
C
Paralog inversion:
E-box Factors
Female
Male
Clock
Npas2
Bhlhe40/Dec1
Bhlhe41/Dec2
FPKM
ZT time (hours)
Daily Expression (FPKM sum)
F
M
D
Other Clock genes
Female
Male
Bmal1
Per1
Per2
Cry1
Cry2
FPKM
ZT time (hours)
Daily Expression (FPKM sum)
F
M
E
Dec2
Dec1
Cry2
Cry1
Per1
Per2
E-box
Dbp
Clock
Npas2
Bmal1
RorC
Revα
Revβ
D-box
RRE
Nfil3
Legend:
Diestrus
Proestrus
Estrus
Metestrus
Day 1
Day 2
Female
Male
* Student's t-test p<0.05
** Student's t-test p<0.01
*** Student's t-test p<0.001
### DoDR meta.p<0.1 males vs DP
#### DoDR meta.p<0.1 males vs EM
##### DoDR meta.p<0.1 males vs DP and EM
###### DoDR meta.p<0.1 DP vs EM

**Figure S5. Core circadian clockwork architecture is altered between males and females.** Gene expression profiles (line plots, some are replotted from Fig 1 for comparison) and daily sum (bar + individual data points) comparisons between male and female clock genes follow three themes: **A.** Shift from activator (*Rorc*) > repressor (*RevErbα/β*) in females to activator < repressors in males, **B** Loop amplitude: activator *Dbp* and repressor *Nfil3/E4bp4* are both more robustly expressed in females compared to males, **C.** Paralog inversion in expression of the E-box transcription factor paralog pairs (*Clock/Npas2* and *Bhlhe40/41*). **D.** Expression profiles of other core clock genes. Differences in overall expression (bar graphs) are denoted by * and differences by DoDR are denoted by # (as described in the legend). **E.** Schematic of core clockwork feedback loops regulating gene expression [2,3], summarizing the differences overall expression/abundance between the sexes: blue = higher expression in males, pink = higher expression in females (grey = no apparent sex differences), green arrow = transcriptional activator, red blunt arrow = repressor. Differences identified by DoDR were not incorporated.

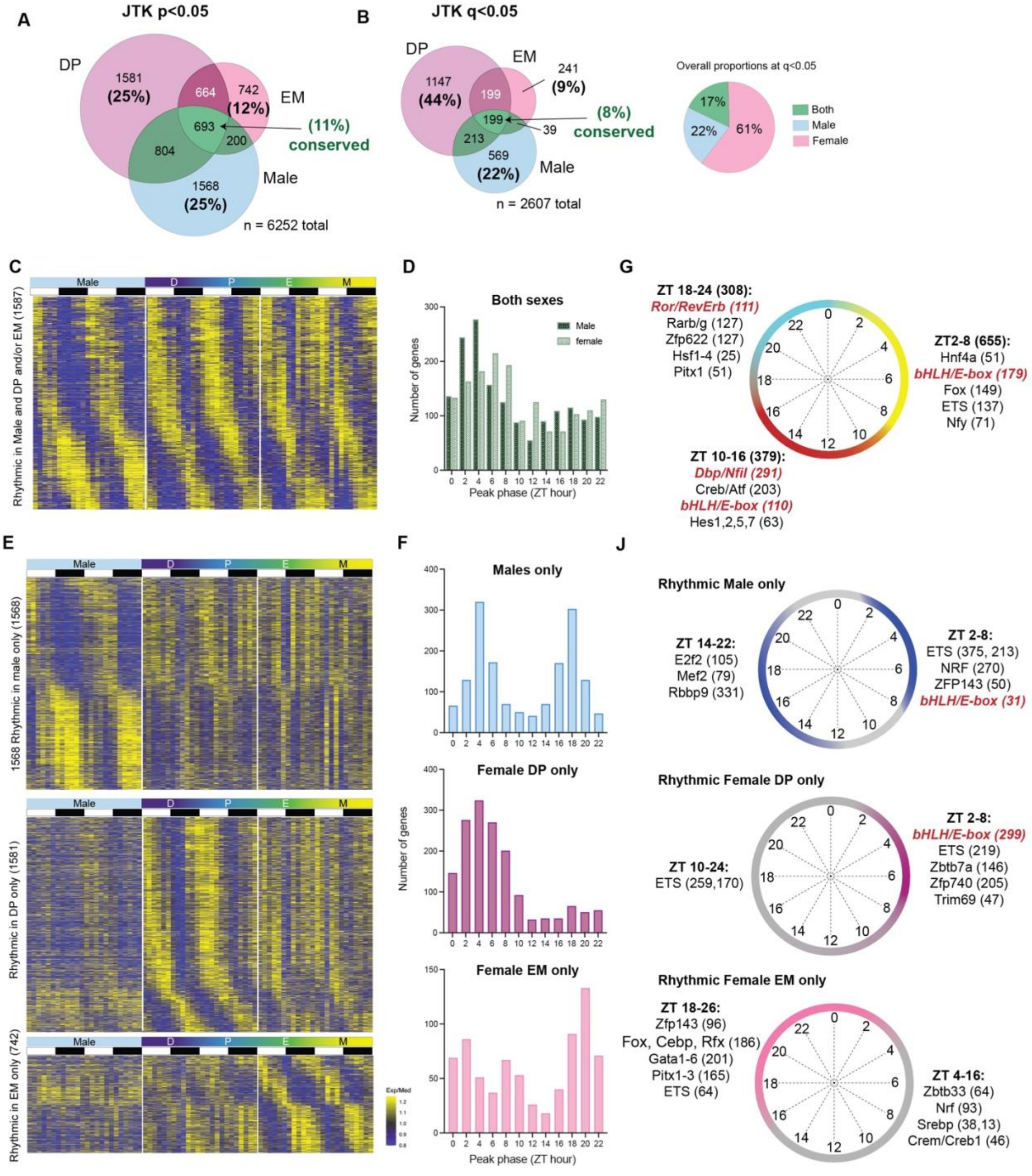


**Figure S6. Estrous cycle expands sex differences in rhythmicity.** Sex-dependent diurnal gene expression in the liver is driven by diverse and complex mechanisms. **A.** Venn diagram comparing the population of rhythmic transcripts (JTK p<0.05) in males and the two rolling estrous cycle stages (DP and EM); the number of transcripts in each section is indicated. See also Fig S4. **B**. Venn diagram comparing the population of rhythmic transcripts (JTK q<0.05) across males, DP, and EM (left) and overall proportions of males and females (right). The number of transcripts in each section is indicated. **C.** Heatmap expression of rhythmic genes across sex. Data (FPKM) were median-normalized across sexes/estrous cycle phases; individual genes run across the entirety of the heat map and are stacked vertically by peak phase. Yellow = high, blue = low. White lines are drawn on top to help separate groups. **D**. Frequency distributions of peak expression phase (ZT hours) for males and females. **E**. Heatmaps of the transcripts for those genes in the non-overlapping groups (from A), plotted as in E. **F.** Frequency distributions of peak phase for the genes in H. **G**. Transcription factor binding site enrichment analysis (iRegulon) of genes peaking at different time intervals across the day in both sexes, along with the top 5 enriched sites (defined by a representative transcription factor or class; numbers of gene targets are shown in brackets). Red highlights the enriched core circadian transcriptional regulators that are canonically responsible for driving peak expression at the indicated times of day. See also Figure S5. **H**. Enriched transcription factor binding sites and potential transcriptional networks for major genes with similar rhythmic expression profiles/peak phases in each group, drawn as in G. Red highlights circadian elements.

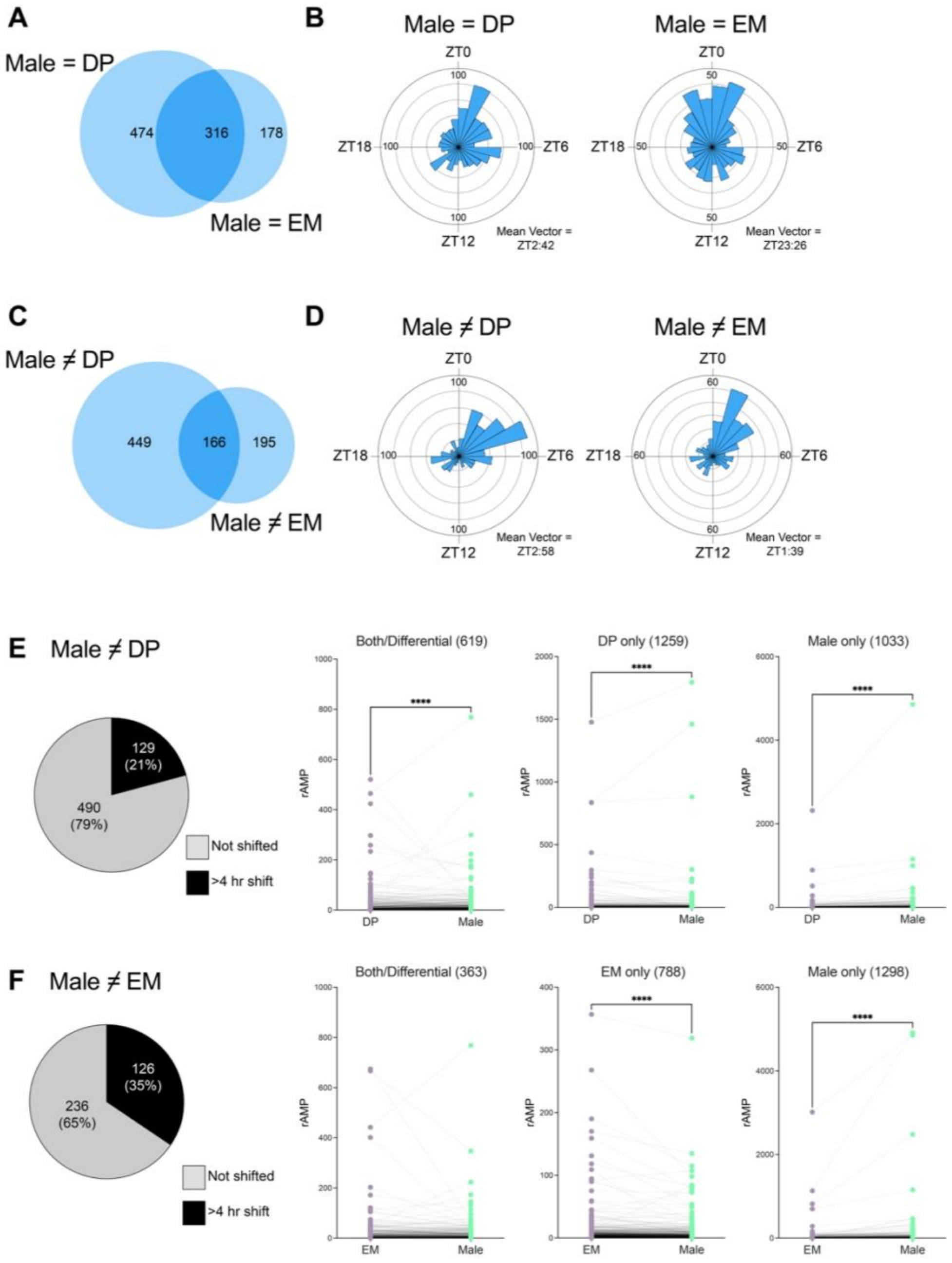


**Figure S7. Differential rhythm characteristics between males and females in DP or EM. A**. Venn diagram of genes considered 'normally' rhythmic ("=" sign) in male vs DP and male vs EM. **B**. Rayleigh plots of the peak phase of the genes in A, the mean vector is shown below. These distributions are significantly different (Watson's $U^2$ test $p<0.001$). **C**. Venn diagram of genes that are differentially rhythmic between male and either DP or EM, but rhythmic in both comparison groups ("≠"). **D.** Rayleigh plots of peak phase of the genes in C, with mean vectors noted. These distributions were also significantly different (Watson's $U^2$ test $p<0.001$). **E-F.** Phase (>4 hours, differential but rhythmic in both) and amplitude differences of differentially rhythmic genes in Male vs.

DP (E) and Male vs. EM (F). Numbers of transcripts in each group are in parentheses. **** = $p<0.0001$, Wilcoxon test.

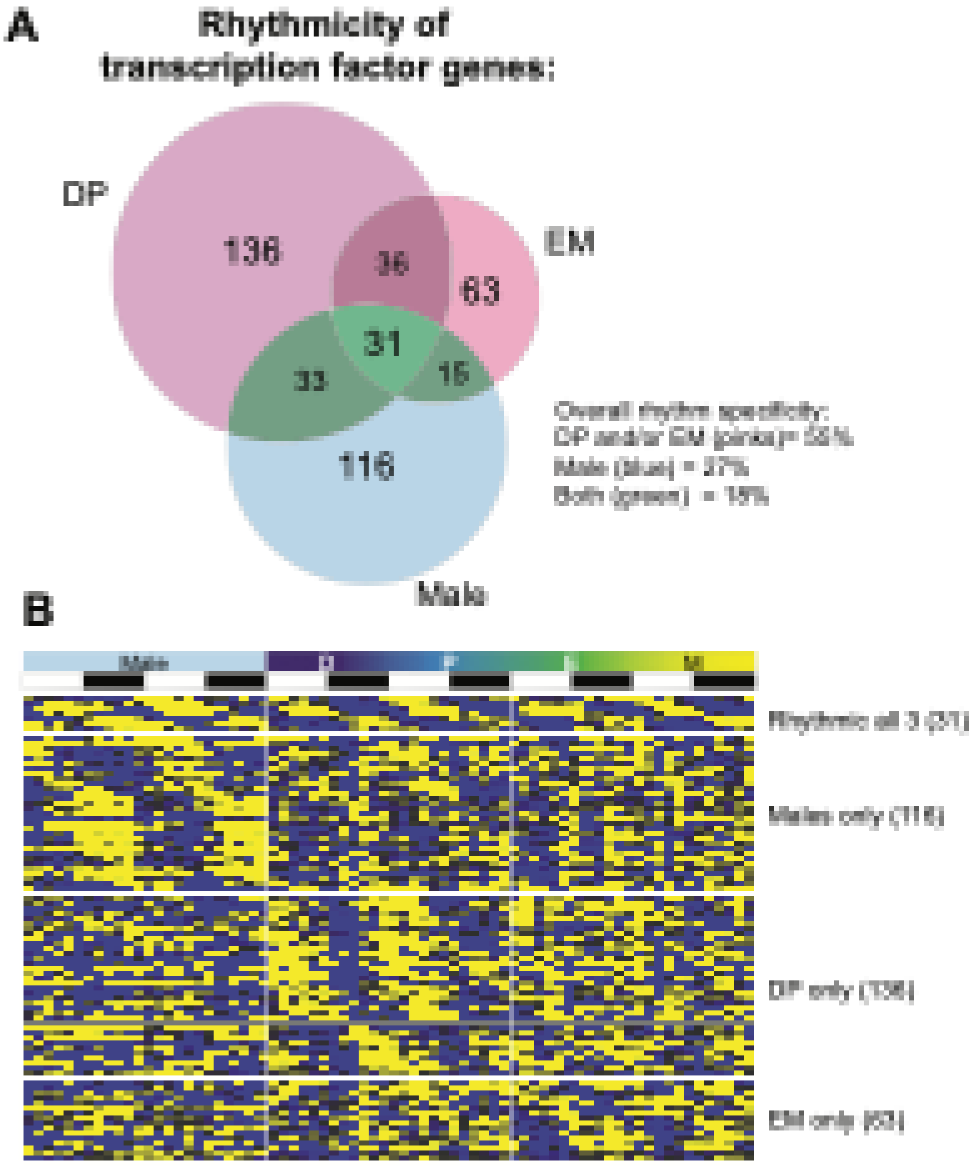


**Figure S8. Differential rhythmicity of transcription factor genes. A.** Venn diagram of rhythmic transcription factor genes in the liver https://cgs.csail.mit.edu/ReprogrammingRecovery/mouse_tf_list.html [4]. **B.** Heatmap for the transcription factors transcripts in the major groups depicted in A.

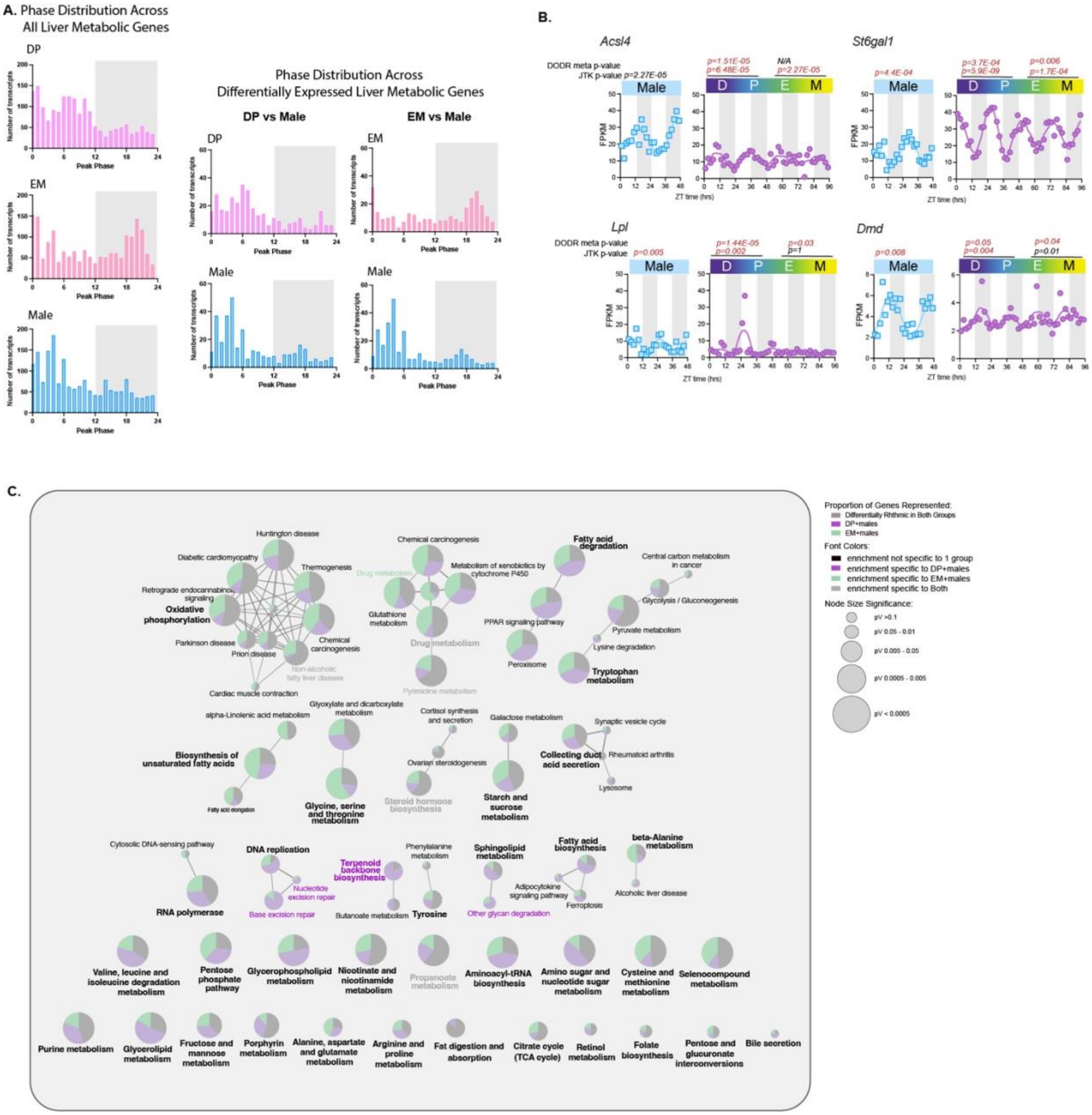

**Figure S9. Comparison of phase distribution and target gene networks of liver metabolic genes. A.** Phase distribution plots of- Left: all liver metabolic genes across males and females (DP and EM) and Right: differentially expressed liver metabolic genes across two groups- DP and males; EM and males [5]. **B.** Examples of 4-day expression profiles of representative genes that change rhythmic expression across sex and e-cycle phase. DoDR and JTK p-values (cut off for significance p<0.05) are indicated above each plot. **C.** Fully annotated KEGG pathway interaction networks of connected signaling pathways for liver metabolic genes [5]. Green (specific to DP vs male comparison), purple (specific to EM vs male comparison), gray (enriched across both comparison groups), and black (not specific) fonts indicate top enriched pathways identified using ClueGO (version 2.5.10) and CluePedia (version 1.5.10) (see Methods).
